# Sonoran Desert *ex situ* conservation gap analysis: charting the path towards conservation

**DOI:** 10.1101/2024.11.06.622087

**Authors:** María Guadalupe Chávez-Hernández, Pablo Gómez-Barreiro, Joseph Douglas Mandla White, Michael Way

**Author notes:** **Corresponding author** María Guadalupe Chávez-Hernández.

## Abstract

Plant biodiversity is under threat. With two out of five plants at risk of extinction, preserving taxa through *ex situ* conservation approaches must be considered a priority. The Sonoran Desert (SD), a highly biodiverse ecoregion shared between Mexico and the United States is home to over 4000 native plant taxa. A method to determine conservation priorities was developed in collaboration with institutions from both countries. Herbaria records and data from nine seed banks were analysed to identify gaps in previous conservation efforts. The SD native taxa were prioritised based on a Priority Score (PS) and their potential distribution was modelled to identify priority regions. 4029 native taxa were reported. 1441 have accessions preserved in seed banks, but only 412 have been collected inside the SD. The PS considers some potentially endemic (126) and threatened taxa (112) as the most urgent to preserve. It also includes information about the storage behaviour (3236 orthodox), useful plants (1406), and taxa without populations in protected areas (230) to categorise the species. Although most of the SD flora is not represented in seed banks, at least 80% is predicted to produce desiccation-tolerant seeds and thus can be cost-effectively stored. Spatial analysis shows that the central region of the Baja California Peninsula stands out for its species richness and endemism. Our study presents the first *ex situ* conservation gap analysis of the SD flora and provides a replicable methodology for identifying priority species and potential areas for *ex situ* collection.

## 1. Introduction

### 1.1. The role of ex situ conservation

Global plant diversity is under threat (Antonelli et al., 2020). The causes are highly diverse, ranging from habitat loss and climate change to the overexploitation or illegal trade of species (Breman et al., 2021; Corlett, 2016; Margulies et al., 2022). It has been estimated that two out of five species of plants might be at risk of extinction, which highlights the urgency of implementing alternatives for their conservation (CBD, 2012). When *in situ* conservation is not viable or complementary protection is required, *ex situ* conservation plays a fundamental role in safeguarding biodiversity (Li and Pritchard, 2009).

Traditional seed banks (low seed moisture content and low storage temperature) are proven cost-effective *ex situ* solutions known to considerably expand the lifespan of most orthodox seeds while maintaining high genetic diversity (Breman et al., 2023). Among them, the Millennium Seed Bank is the largest repository of wild plant species, banking more than 2.4 billion seeds from almost 40,000 species (Breman et al., 2021; RBG Kew, 2023). Its collaboration since 2000 with institutions in around 100 countries or territories stands out, consolidating the Millennium Seed Bank Partnership (MSBP) as the world”s largest ex situ conservation programme (Liu et al., 2018). Despite the importance of its collections and standardization of its associated data, the representativeness of specific priority groups including endangered species, useful plants, or entire regional floras in MSBP seed banks has usually been evaluated individually, for example, Godefroid et al. (2011), Ramírez-Villegas et al. (2010), Rivière and Müller, (2018), and Teixido et al. (2017).

### 1.2. Prioritisation within the Sonoran Desert ecoregion

We identified that gap analyses are a useful methodology to identify opportunity areas, propose priorities, and improve future conservation efforts, which is especially important in highly diverse and threatened regions (Carver et al., 2021; Scott et al., 1993; Sowa et al., 2007). The Sonoran Desert (SD), located in Mexico and the United States, meets these criteria, making it a relevant study area where evidence of gaps can help prioritise ecoregional conservation actions. The SD is the hottest and most diverse desert in North America, including a wide variety of plant life forms, from columnar cacti and woody shrubs to winter and summer annual herbs (Burquez et al., 1999; Rebman and Roberts, 2012; Robichaux, 1999; Weiss and Overpeck, 2005). The environmental variation, rainfall regimes, and community composition are such that at least seven subdivisions have been proposed to differentiate specific areas within the SD (Shreve and Wiggins, 1951; Turner and Brown, 1982). Regarding specific diversity, calculations estimate that between 3,200 and 3,400 species of plants occur in its territory (McLaughlin and Bowers, 1999). Amongst this diversity, keystone species such as the saguaro cactus (*Carnegiea gigantea*) or unique parasitic plants like *Pholisma sonorae* (Fig. 1B and E) stand out for their environmental and cultural relevance (Drezner, 2014).

**Fig. 1.**
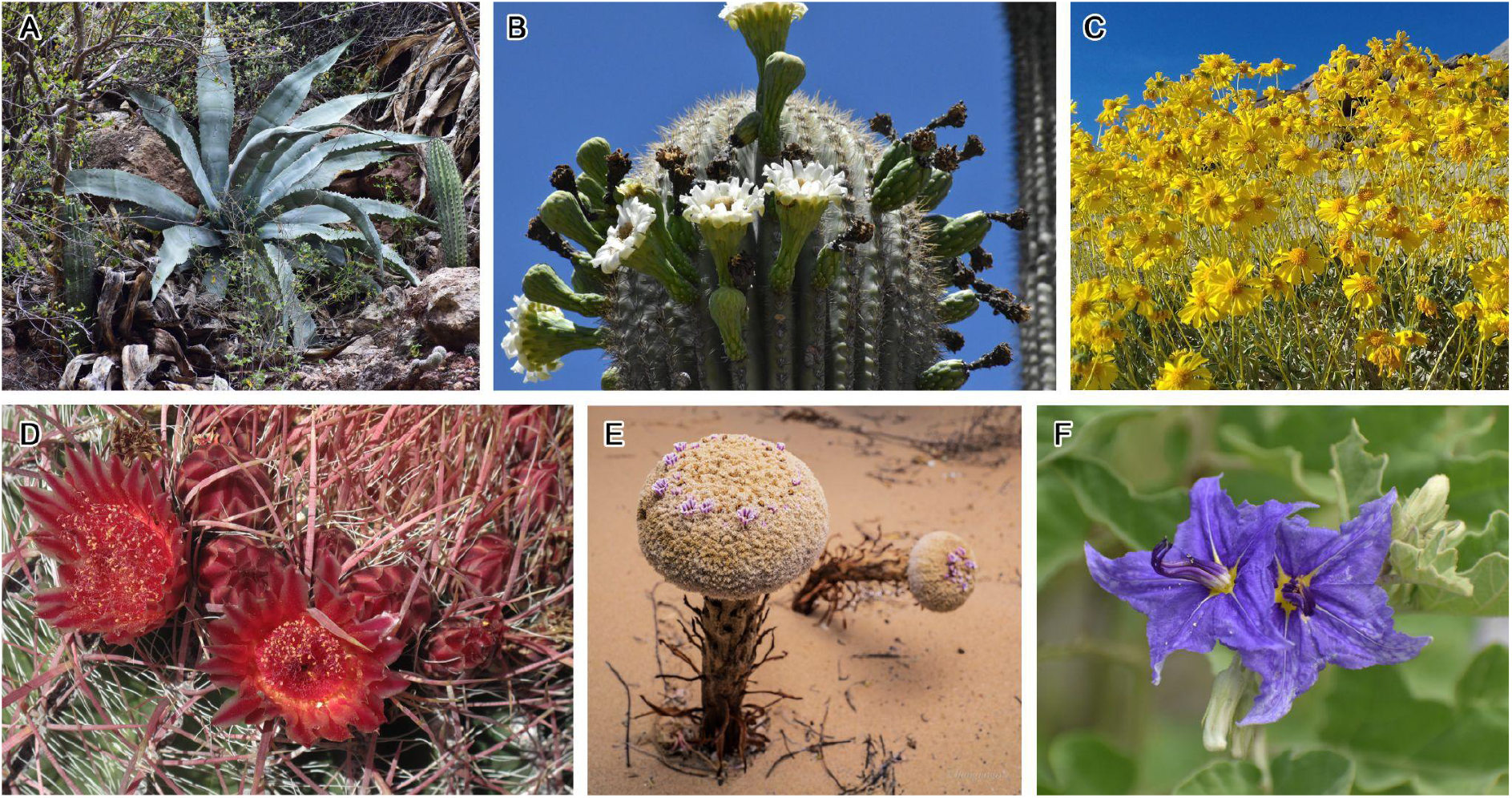
Examples of the Sonoran Desert flora. (A) *Agave colorata* Gentry. (B) *Carnegiea gigantea* (Engelm.) Britton & Rose. (C) *Encelia farinosa* A. Gray ex Torr. (D) *Ferocactus gracilis* H.E. Gates. (E*) Pholisma sonorae* (Torr.) Yatsk. (F) *Solanum houstonii* Martyn. Images by Armando Ponce-Vargas (A, B, C, F), Wolfgang Stuppy (D) and Juan Diego Arias-Montiel (E).

However, this biodiversity faces threats such as climate change and the spread of invasive species (Tinoco-Ojanguren et al., 2016; Weiss and Overpeck, 2005; Wilson et al., 2002). Recent studies on annual plants reported a potential increase in the species” extinction risk if rainfall regimes continue changing (Cuello et al., 2022). Also, analyses of specific taxa, such as members of the Cactaceae family, have shown the potential decline of their populations in the face of climate change (Breslin et al., 2020). Some modifications in the limits of the SD have also been predicted, so the distribution of its species is likely to change in the future (Weiss and Overpeck, 2005). It is important to note that most of the flora from arid and highly seasonal environments, such as the SD, is usually associated with orthodox seed behaviour (Tweddle et al., 2003; Wyse and Dickie, 2017). Such evidence suggests that the SD flora could be primarily stored in seed banks in future ex situ conservation actions.

This study is focused on the analysis of prior knowledge of the SD flora, using compiled herbaria records and seed bank data to determine the gaps in previous collection efforts. Priority taxa are suggested based on a Priority Score (PS), which was calculated from six individual scores, and priority areas for *ex situ* conservation were identified using presence records and species distribution models (SDM).

## 2. Materials and methods

### 2.1. Site delimitation and data collection

The SD area includes the Sonoran Desert and Baja Californian Desert ecoregions of the Level III Ecoregions of North America (EPA, 2023). This approach corresponds to previous delimitations for the SD (Shreve and Wiggins, 1951; Turner and Brown, 1982) and the phytogeographic region classification of the Baja California Peninsula proposed by Gonzalez-Abraham et al. (2010).

Presence records were downloaded from the consortium of herbaria SEINet (2023) selecting the quadrant 42.2°N, 22.5°N, 124.5°W, 108.2°W, which includes the states of Arizona and California in the United States and Baja California, Baja California Sur, and Sonora in Mexico. Hidden records were requested from the Symbiota Portal Hub to include distribution data for protected species. The complementary seed bank accession data were compiled from the MSBP Data Warehouse, and seven associated collaborators (Table 1). All records (N=1,045,471) were cleaned by removing duplicates and deleting records without coordinates, their taxonomy was matched against The World Checklist of Vascular Plants, and the native SD species were selected using the information available in Plants of the World Online (Govaerts et al., 2021). After cleaning and name matching, there were 1271 species with one to two presence records, 1067 species with three to ten, and 1691 species with more than ten records.

**Table 1.** Seed data providers, number of accessions, and taxa collected for each institution. The total number of accessions is reported.

| Institution | Abbreviation | Total number of seed accessions | Total number of native SD taxa preserved | Number of accessions in the SD | Number of native SD taxa preserved in the SD |
| --- | --- | --- | --- | --- | --- |
| California Botanic Garden | CalBG | 2188 | 766 | 740 | 248 |
| Facultad de Estudios Superiores de Iztacala | FESI | 455 | 342 | 244 | 160 |
| Millennium Seed Bank | MSB | 1881 | 1028 | 666 | 289 |
| Santa Barbara Botanic Garden | SBBG | 16 | 10 | 0 | 0 |
| San Diego Botanic Garden | SDBG | 28 | 21 | 5 | 5 |
| San Diego Zoo Wildlife Alliance | SDZWA | 319 | 190 | 106 | 53 |
| University of California Santa Cruz | UCSC | 216 | 65 | 6 | 4 |
| The Arboretum at Flagstaff | thearb | 80 | 36 | 10 | 2 |
| **Total** |  | **5183** |  | **1777** |  |

### 2.2. Species prioritisation

Additional data were obtained and analysed for all the native SD taxa. Six individual scores were calculated for each of the species: 1) Endemism Score (EndS), 2) Conservation Status Score (ConsStS), 3) Storage Capacity Score (StorS), 4) Useful Plants Score (UsefS), 5) In-situ Conservation Score (InSituS), and 6) Seed Bank Accessions Score (SBS). The possible values for those scores are specified in Table 2.

**Table 2.** Possible values for each taxon in the six individual scores of the Priority Score (PS).

| Individual score and possible categories | Value |
| --- | --- |
| 1. Endemism Score (EndS) |  |
| a. Not endemic to the SD | 0 |
| b. Endemic to the SD | 1 |
| 2. Conservation Status Score (ConsStS) |  |
| a. IUCN Red List |  |
| i. Not evaluated taxa | 0 |
| ii. Data Deficient (DD) | 0.5 |
| iii. Least Concern (LC) | 0 |
| iv. Near Threatened (NT) | 0.25 |
| v. Vulnerable (VU) | 0.5 |
| vi. Endangered (EN) | 0.75 |
| vii. Critically Endangered (CR) | 1 |
| b. NOM-059-SEMARNAT-2010 |  |
| i. Not evaluated taxa | 0 |
| ii. Sujeta a protección especial (Subject to special protection) (Pr) | 0.33 |
| iii. Amenazada (Threatened) (A) | 0.66 |
| iv. En peligro de extinción (Endangered) (P) | 1 |
| c. The United States Endangered Species Act (ESA) |  |
| i. Not evaluated taxa | 0 |
| ii. Threatened | 0.5 |
| iii. Endangered | 1 |
| 3. Storage Capacity Score (StorS) | 1-probability of recalcitrance given by the tool |
| 4. Useful Plants Score (UsefS) | (# Uses per spp / 10 (maximum)) |
| 5. In-situ Conservation Score (InSituS) |  |
| a. At least one population inside a PA in the SD | 0 |
| b. At least one population inside a PA outside the SD | 0.5 |
| c. Not populations inside PA | 1 |
| 6. Seed Bank Accessions Score (SBS) |  |
| a. High or medium-high usability accessions in the SD | 0 |
| b. High or medium-high usability accessions outside the SD | 0.25 |
| c. Low or medium-low usability accessions in the SD | 0.5 |
| d. Low or medium-low usability accessions outside the SD | 0.75 |
| e. No accessions in seed banks | 1 |

A Priority Score (PS) between 0 and 1 was calculated using the formula:

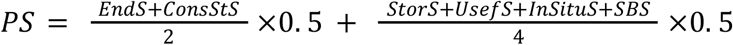

With this approach, the potentially endemic and threatened taxa are given double the weighting over other scores to align the PS with the goals of this project and Target 8 of the Global Strategy for Plant Conservation, which is focussed on the inclusion of endangered species in *ex situ* programmes (CBD, 2012). The taxa with values closer to 1 would have a higher level of priority.

#### 2.2.1. Endemism score (EndS)

The native taxa list was used as a starting point for the selection of potentially endemic species. Those occurring outside of the delimited SD area were filtered out.

#### 2.2.2. Conservation status score (ConsStS)

The conservation status score of the taxa was assessed using three sources of information, including both international and regional assessments. Information from the IUCN Red List of Threatened Species (IUCN, 2023) was extracted. Because there is evidence that “Data Deficient” taxa likely include a significant number of threatened species, they received a value of 0.5 (Borgelt et al., 2022; Parsons, 2016).

Mexican regional assessments were recovered from the “Norma Oficial Mexicana NOM-059-SEMARNAT-2010” (SEMARNAT, 2010). United States regional assessments from the United States Endangered Species Act were obtained from the NatureServe Explorer (NatureServe, 2023).

#### 2.2.3. Storage Capacity Score (StorS)

The probability of recalcitrance of the species was calculated using the Seed Storage Behavior Predictor tool (Wyse and Dickie, 2018). The score was obtained by subtracting the predicted value from one, prioritising those taxa whose seeds are predicted to have orthodox i.e. storable behaviour.

#### 2.2.4. Useful Plants Score (UsefS)

The number of reported uses per species was obtained from the World Checklist of Useful Plant Species Database. This was divided by the maximum number of uses (10): medicines, materials, environmental uses, human food, gene sources, animal food, poisons, social uses, fuels, and invertebrate food (Diazgranados et al., 2020).

#### 2.2.5. In-situ Conservation Score (InSituS)

Protected Areas (PA) shapefiles were downloaded from Protected Planet: The World Database on Protected Areas (UNEP-WCMC & IUCN, 2023). The presence of populations within protected areas, both inside and outside the SD, was evaluated.

#### 2.2.6. Seed Bank Accessions Score (SBS)

The presence and usability of native taxa accessions were analysed separately: 1) inside the SD and 2) outside the SD. This distinction was made since accessions in the SD have more relevance to the aims of this study, and even though some taxa might have backup collections in other regions, their SD populations should also be preserved.

The usability of the collections was evaluated using five individual component scores modified from previous gap analyses (Dhanjal-Adams, et al., Unpublished results): 1) Accessions Score (AccS), totalling the number of accessions per taxon, regardless of its institution of origin. 2) Seed Count Score (SCS), which evaluates the presence or absence of a seed count (either a current count or an adjusted count). 3) Seed Number Score (SNS), modified from previously recommended target seed numbers (Way, 2003). 4) Germination Test Score (GerTestS), which evaluates the presence or absence of a germination test, and 5) Viability Score (ViabS), defined as the value of the reported viability divided by 100. The possible values given for each taxon are specified in Table 3.

**Table 3.** Possible values for each taxon in the five individual scores of the Seed Usability Score (SeedUsS).

| Individual score category | Value |
| --- | --- |
| 1. Accessions Score (AccS) |  |
| a. 1 | 0 |
| b. 2-5 | 0.2 |
| c. 6-9 | 0.4 |
| d. 10-19 | 0.6 |
| e. 20-29 | 0.8 |
| f. ≥30 | 1 |
| 2. Seed Count Score (SCS) |  |
| a. No | 0 |
| b. Yes | 1 |
| 3. Seed Number Score (SNS) |  |
| a. <500 | 0 |
| b. 500-1000 | 0.2 |
| c. 1000-5000 | 0.4 |
| d. 5000-10000 | 0.6 |
| e. 10000-20000 | 0.8 |
| f. >20000 | 1 |
| 4. Germination Test Score (GerTestS) |  |
| a. No | 0 |
| b. Yes | 1 |
| 5. Viability Score (ViabS) | Viability/100 |

A Seed Accessions Usability Score (SeedUsS) between 0 and 1 was calculated using the formula:

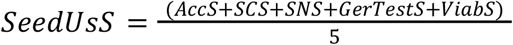

The taxa with values closer to 1 have higher usability in their collections and, therefore, less priority in the PS. Four usability categories were proposed: 1) High usability (SeedUsS >= 0.75); 2) Medium-high usability (SeedUsS >= 0.5 < 0.75); 3) Medium-low usability (SeedUsS >= 0.25 < 0.5); and 4) Low usability (SeedUsS < 0.25).

### 2.3. Geographical prioritisation

Collection density maps using a logarithmic scale were created to visualise the sampling effort of both data sources: herbarium collections and seed bank accessions. To obtain priority heatmaps, species distribution models (SDMs) were generated when the taxa had more than ten records. For the taxa with between three and ten records, geographical models (GMs) using an inverse distance weighting approach were produced. For the taxa with fewer than three records, buffered points with a radius of 10 km were created around the presence coordinates (Brown et al., 2014; Tovar et al., 2023). SDM, GM, and buffer maps were then summed to obtain overall collection heatmaps of priority regions.

For the SDMs, the nineteen bioclimatic variables from WorldClim version 2.1 (Fick and Hijmans, 2017) were considered. A Pearson correlation analysis with a threshold of 80% was carried out to reduce collinearity between the environmental variables. Eight non-correlated variables were selected, primarily covering temperature and precipitation variables (Yin et al., 2022). To reduce spatial bias in the occurrence records, we spatially thinned presence records by randomly choosing one record for each +-1 km^2^ pixel, using our environmental layers as a reference grid. After that, 10,000 pseudo-absences were randomly created for each species. Pseudo-absences were constrained to only occur outside of a 10 km buffer of a species” presence records.

To model species distributions, we used the well-known Maxent distribution model v 3.4.1 (Phillips et al., 2006), implemented in the SDMtune R package (Vignali et al., 2020). Models were fitted with an automatic selection of feature responses, including linear, quadratic, product, and hinger feature classes. Each species β regularization coefficient went through a tuning process to control model complexity and reduce potential overfitting. The models were fitted with a range of regularization values: 0.25, 0.5, 1, 2, 4, and 8 (Radosavljevic and Anderson, 2014). In addition, models were fitted using spatial cross-validation, where presence records were split into four groups of equal (or as close to as possible) size, by first finding the latitude line that splits the data into two equal groups, followed by the longitude line (Kass et al., 2021). This produces four spatial folds. The model is then iteratively run using only 3 folds and tested on the remaining spatial fold. Each model”s performance was evaluated using the area under the curve of the receiver operator characteristic plot (AUC) value. Models with the highest AUC value for each respective regularization coefficient were selected for each species and the mean AUC was calculated across each spatial fold iteration. Species with an AUC value below 0.7 were removed from the downstream analysis, while for the remaining species, we predicted their distributions across the SD area against the chosen environmental variables.

For species with fewer presence records than environmental variables (i.e. between 3 and 9 presence records), we used a form of geographical models (GMs) using inverse-distance weighting (Pironon et al., 2024). We first created pseudo-absence records using the same protocol as the one used with the SDMs. Models were evaluated using a leave-one-out approach, where we split the dataset into k-folds based on the number of presence records for each species. For each fold, we iteratively produced a GM based on n-1 presence records and a ratio of n-1/n pseudo-absences and evaluated the models on the withheld presence record and pseudo-absence records. We calculated the mean AUC value across all model iterations and treated species with AUC values below 0.7 in the same way as species with fewer than three occurrences. The final distribution was the mean map prediction of each GM iteration.

For species with fewer than three presence records, we produced a 10 km radial buffer around each record and directly rasterized this into a binary output across the SD area, with regions within the buffers classed as 1s and outside as 0s. To identify regions for prioritisation, heatmaps were generated by stacking the generated probability (SDMs and GMs) and binary (buffer) maps. This is analogous to the approach used to determine species richness in stacked-species distribution models (S-SDMs). Heatmaps were generated by 1) using the 100 taxa with the highest PS (12 SDMs + 45 GMs + 41 buffered areas), 2) considering potentially endemic taxa (17 SDMs + 53 GMs + 54 buffered areas), and 3) modelling the distribution for threatened taxa (63 SDMs + 34 GMs + 15 buffered areas). Climatic data could not be extracted from two species (*Eriogonum angelense* and *Martynia palmeri*), so they were not added to the analysis.

### 2.4 Data analysis

All data cleaning, visualisation and analyses in this paper were conducted using R (version 4.2.3) (R Core Team, 2023) and the following packages: *corrplot* (Wei et al., 2017), *dismo* (Hijmans et al., 2017), *ENMeval* (Kass et al., 2021), *flexsdm* (Velazco et al., 2022), *SDMtune* (Vignali et al., 2020), *sf* (Pebesma 2018), *terra* (Hijmans et al., 2022), *tidyverse* (Wickham et al., 2019), and *virtualspecies* (Leroy et al., 2016)

### 2.5 Data availability

The majority of the botanical occurrence data is freely available from the consortium of herbaria SEINet. Unpublished Seed Bank accession data and our compiled georeferenced data cannot be shared due to the confidentiality of the locations of many of the endangered species. For access to these data, please refer to the sources cited in the Materials and Methods section and contact sources directly for requests.

## 3. Results

### 3.1. Status of the Sonoran Desert flora

4029 native taxa (species, subspecies, varieties, and hybrids) from 149 families were identified for the SD (Fig. 2). The families with the largest number of taxa were Asteraceae (611), Fabaceae (410), and Poaceae (249).

**Fig. 2.**
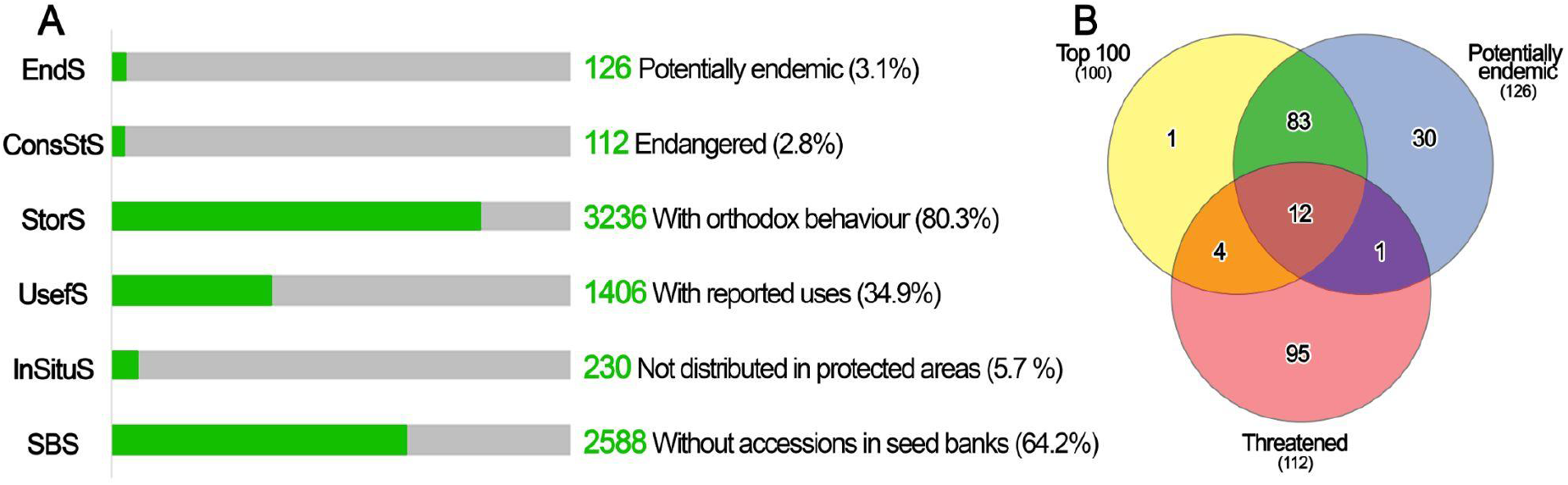
Status of the 4029 taxa recorded for the Sonoran Desert according to the six individual scores, the proportion of taxa prioritised by the scores is shown in green (A). Venn Diagram showing the proportion of shared taxa according to the top 100, potentially endemic and threatened lists (B).

#### 3.1.1. Endemism

126 taxa from 35 families were identified as potentially endemic from the SD (Fig. 2A). The most diverse families were Cactaceae (18), Asteraceae (15), and Polygonaceae (11). Eight of the potentially endemic species have been seed banked and only one of those has high usability accessions inside the SD (Fig. 3A, B).

**Fig. 3.**
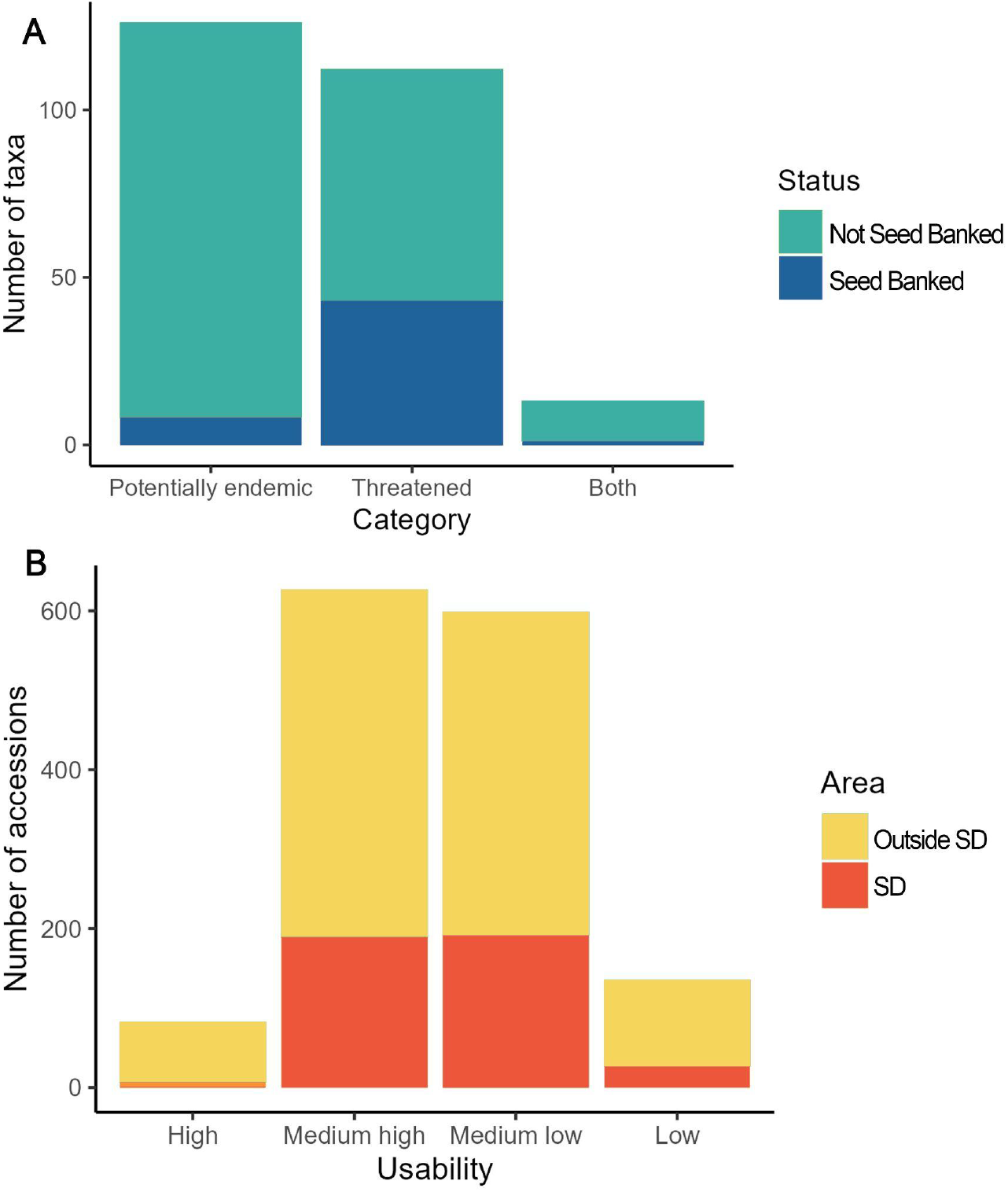
**A**. Status of the potentially endemic and threatened taxa in seed banks. **B**. Usability of the seed accessions of native SD taxa considering the values of the Seed Usability Score.

#### 3.1.2. Conservation status

112 taxa are considered under threat according to the three studied data sources (Fig. 2A). 644 taxa have been assessed by the IUCN. 568 were defined as “Least Concern” and 16 were considered “Nearly Threatened”. 10 of the taxa were classified as “Data Deficient”. 50 taxa were considered threatened: 31 as “Vulnerable”, 17 as “Endangered”, and 2 as “Critically Endangered”. 55 taxa are listed in the NOM-059-SEMARNAT-2010. 36 of them as “subject to special protection”, 14 as “threatened”, and 5 as “endangered”. Five taxa were listed as endangered under the United States Endangered Species Act. Only 43 of the 112 threatened taxa have been seed banked, 19 of them inside the SD (Fig. 3A).

#### 3.1.3. Storage Capacity Score

3177 taxa were predicted to have orthodox seed behaviour (80%). Only 34 were classified as recalcitrant and belonged mainly to the genus *Quercus* (22 taxa). There was not enough information to assign the potential behaviour of 818 taxa.

#### 3.1.4. Useful Plants

1406 taxa are included in the World Checklist of Useful Plant Species with at least one use. 137 of them have five or more reported uses. In this study, all the uses were considered to have the same importance.

#### 3.1.5. In-situ Conservation

230 taxa do not have any populations inside protected areas. 989 taxa have at least one record inside a protected area outside the SD polygon, and 2810 are distributed in protected areas inside the SD.

#### 3.1.6. Seed Bank accessions

5183 accessions from 1441 native SD taxa have been preserved in eight seed banks, but only 1777 accessions from 412 taxa have been collected inside the SD polygon (Table 1). 101 of those 412 taxa have more than five accessions. According to the SeedUsS, 6 of the 412 collected taxa inside the SD have high usability accessions, 189 reported medium-high usability, 191 medium-low usability, and 26 low usability. On the other hand, 76 of the 1029 taxa outside the SD had high usability accessions, 437 medium-high usability, 407 medium-low usability, and 109 low usability (Fig. 3B).

### 3.2. Species prioritisation

Native SD taxa were ranked according to their PS. The top 100 was selected as a priority for seed banking (Supplementary material). The families with most included taxa were Cactaceae (14), Asteraceae (13), Polygonaceae (8), Asparagaceae, Fabaceae, and Rubiaceae (6). 95 potentially endemic and 16 endangered taxa are included in the selection. The genus *Agave*, specifically the species *A. pelona, A. turneri*, and *A. zebra*, were identified as the top priority taxa.

### 3.3. Geographical prioritisation

The density of herbarium collections is higher than the density of seed bank accessions in the SD (Fig. 4). Likewise, the collection bias in the United States portion of the region is evident but differs according to the data source. While herbarium collections are concentrated in Arizona, seed bank accessions are skewed towards California.

**Fig. 4.**
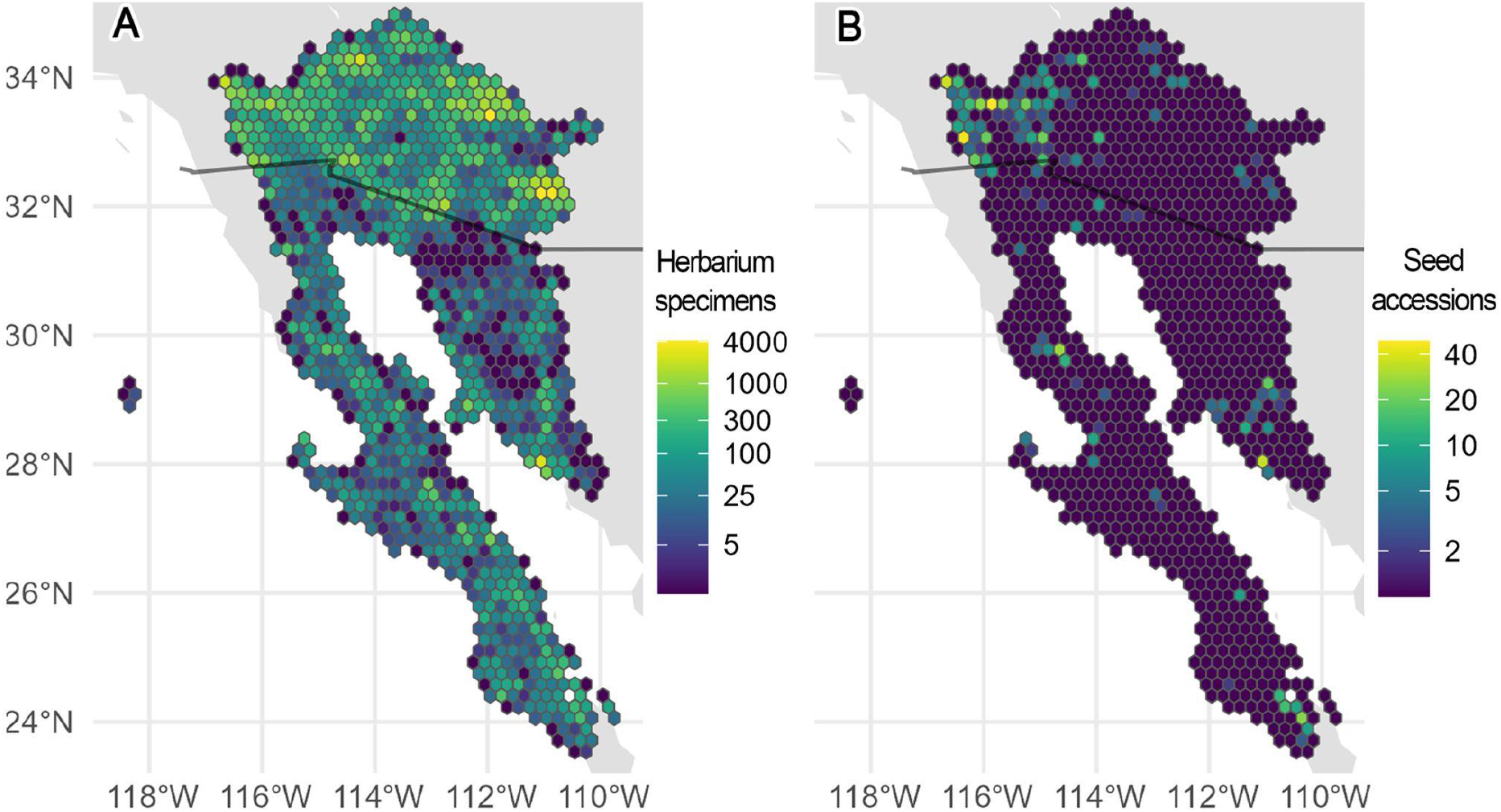
Density of collections from herbaria (A) and seed banks (B) in the Sonoran Desert. The Mexico-US border is highlighted in black. A logarithmic transformation was applied to the data.

The area under the curve (AUC) for all the models was at least 0.7, and 83% was greater than 0.9. Therefore, we consider that the models are acceptable based on previous studies (de Souza and De Marco, 2014; Girardello et al., 2009).

According to the heat map for the 100 taxa with the highest PS, the Baja California Peninsula is where the greatest diversity of priority taxa is distributed (Fig. 5A). The Vizcaino Desert and the northern part of the Central Gulf Coast stand out, including the Ángel Guardían island in the Gulf of California and the Cedros island in the Pacific Coast. Because the PS gives more value to the potentially endemic taxa, the prediction map for those species (Fig. 5B) is similar to the top 100 prediction, sharing 95 taxa between the two predictions (Fig. 2B). The west coast of the Baja California Peninsula is highlighted again as a zone of relevance and Cedros island is predicted to have adequate environmental conditions for at least sixteen taxa, making it a promising area for future seed collection. When considering threatened taxa, priority areas are mostly located in the state of Baja California Sur. The southeast part of the Central Gulf Coast and the Magdalena Plains stand out in the analysis, as well as the southern region of the SD in the state of Sonora. The Cedros island is highlighted again on this map.

**Fig. 5.**
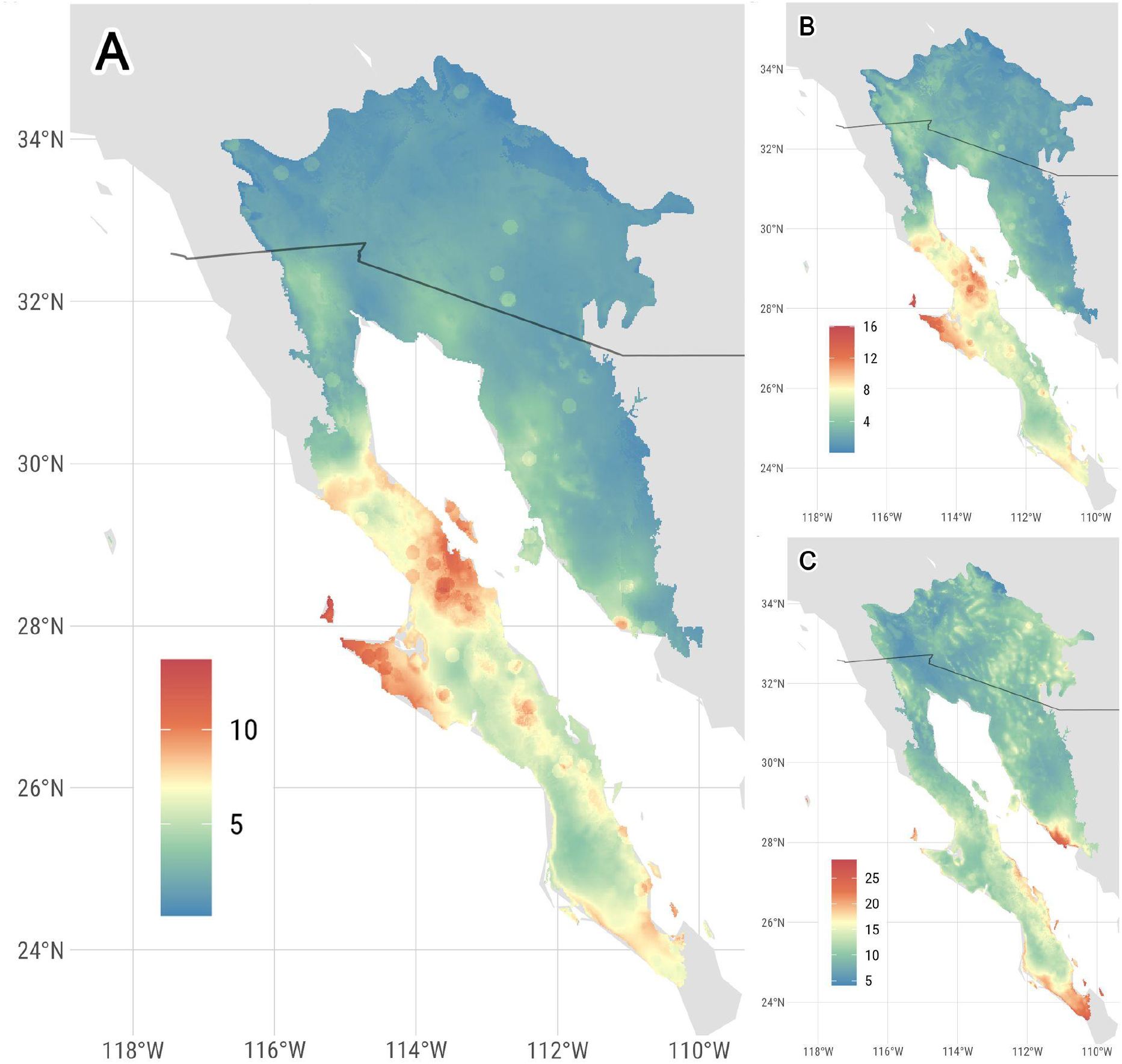
Geographical prioritisation for future seed collections in the Sonoran Desert. (A) Top 100 taxa with the highest Priority Score. (B) Potentially endemic taxa (N=126). (C) Threatened taxa (N=112). The color gradient indicates the number of taxa for which the specific region has suitable climatic conditions for their potential distribution.

## 4. Discussion

### 4.1. SD flora and its representativeness in seed banks

The results of this study indicate that increasing the collection effort is essential to ensure the representativeness of the SD flora in seed banks. Nearly 35% of the native taxa have been seed banked (Fig. 2A), but only 10% of the total number of species have been collected inside the SD area (Table 1). Even though the arid zones of Mexico were mentioned as priority areas for the collection and storage of seeds two decades ago (van Slageren, 2003) there is still a lack of accessions in most of the Mexican part of the SD (Fig. 4B). The number of projects and institutions focused on the study of desert flora, as well as the time they have been in operation (for example, the Sonoran Desert Conservation Plan or the CPC”s Partnership with the California Plant Rescue) could explain the collection bias in the United States, particularly in the state of California (Fig. 4B).

Although most of the SD flora is not represented in seed banks, at least 80% is predicted to produce desiccation-tolerant seeds and thus can be cost-effectively stored in seed banks (Fig. 2A). This pattern has been reported for drylands and it has been proposed that most arid zone species have intrinsic mechanisms that allow them to withstand low humidity conditions (Li and Pritchard, 2009). For species that cannot be seed banked, alternatives such as cryopreservation and propagation in botanic gardens should be considered for their *ex situ* conservation (Pammenter and Berjak, 2013; Pence, 2013; Pence et al., 2020).

Seed collections should contain adequate material to conduct germination and longevity research, as well as represent a valuable source of germplasm for reforestation projects (Breman et al., 2021; Merritt and Dixon, 2011). To achieve that, it is important to have either single high-usability accessions or multiple separate accessions for each taxon. In this study, only 1.46% of the seed-banked taxa in the SD have high usability accessions (Fig. 3B) and only 101 (2.5%) taxa have been collected from more than five seeding populations in the SD. Besides usability, five is also the minimum number of collections likely to ensure a representative sample of the genetic diversity of the species (Liu et al., 2023), which highlights the relevance of carrying out re-samplings of some key species.

### 4.2. Species prioritisation

To reduce the *ex situ* conservation gap, priority species should be determined according to the project”s goals and the available resources. In this work, endemic and threatened taxa were considered to have more relevance in the PS. Previous proposals highlight the relevance of taxa with restricted distribution, which most of the time are also under threat (Burlakova et al., 2011; Godefroid et al., 2011; Kraus et al., 2023). On the other hand, Target 8 of the Global Strategy for Plant Conservation mentions that having 75% of threatened species protected in seed banks, preferably in the country of origin, is essential to ensure their adequate representation in conservation efforts (CBD, 2012).

In the SD, only 6.3% of the potentially endemic taxa and 38.4% of the threatened species have been safeguarded. Countries such as Australia have reported that 84% of endemic flora and 67.7% of threatened species are already seed-banked (Martyn-Yenson et al., 2021). As another example, almost 70% of the European flora and 27% or 44% of its species (depending on the threatened list considered) have backup collections in seed banks (Godefroid et al., 2011). The alarmingly low values for the SD flora only reinforce the idea of proposing new seed collection projects focused on these two large groups of taxa.

We propose a top 100 taxa (Supplementary material) based on the PS values. Cactaceae stands out as the family with the highest number of taxa, which is relevant because there is evidence of both, the importance of this family in the composition and functioning of arid zones and the great threats faced by its species (Hultine et al., 2023, 2016; Ortega-Baes and Godínez-Alvarez, 2006). The genus *Agave* is also considered a priority, with three species included in the top 10. Their environmental and social relevance has been widely reported, mostly because of their close relationship with bats and their traditional use to produce alcoholic beverages (Burke et al., 2019; Trejo-Salazar et al., 2016). Also, most of the *Agave* species are included in the IUCN Red List (Alducin-Martínez et al., 2023).

Nevertheless, additional studies should be performed on the endemic SD taxa, especially to provide a definitive list of endemism and threatened status. An intensification of the sampling of these species to understand the status of their populations and their geographical distribution is also necessary. Since most of the plant species have not yet been assessed in the IUCN Red List (Bachman et al., 2018) and only 644 of 4029 SD species have published assessments, studies of the threatened status of the SD species should be considered a priority. In the last few years, the IUCN Sonoran Desert Plant Specialist Group has played an important role in this mission (Rowe and Montijo, 2020), but there is still a great opportunity area to improve the conservation actions for the SD flora.

### 4.3. Geographical prioritisation

The spatial analyses of this project show that there is a greater abundance of herbarium collections over seed bank accessions (Fig. 4). In addition, seed collections are concentrated in specific areas, mainly in the United States, the vicinity of Hermosillo, Sonora, and the southern region of SD in Baja California Sur (Fig. 4B). Since previously seed-banked taxa have less priority according to the PS, some of these areas might not correspond to the priority zones predicted by the SDMs (Fig. 5). Previous gap analyses in China report similar disparities between the collection zones and location of *ex situ* conservation facilities and the areas of greatest species richness or conservation priorities, often driven by socioeconomic factors (Ye et al., 2023). Considering these factors can help to understand the biases present in the SD as well as propose new collection efforts in the future.

Although those collection biases can also influence the results of spatial analyses, such as the prediction of suitable areas, they are one of the best alternatives to proposing priority conservation zones (Hughes et al., 2021). For this project, most of the endemic or threatened taxa did not have at least ten presence records to generate informative SDMs, so buffered areas and GMs were created to include the information of all the taxa. Still, this indicates that carrying out botanic sampling directed at these groups of plants remains to be done.

The predicted priority areas for the 100 taxa with the highest Priority Score are mainly located in the Baja California Peninsula (Fig. 5A). This region has been reported to be highly diverse, with at least 4000 species of plants and 30% endemism (Rebman and Roberts, 2012; Wiggins, 1980). This diversity and endemism could be explained by the Peninsula”s isolation and its environmental heterogeneity (Riemann and Ezcurra, 2005). It also explains why almost the same areas are predicted as a priority for potentially endemic taxa (Fig. 5B). According to the threatened taxa list, the areas where environmental variables are most suitable for these species are mainly found in Baja California Sur (Fig. 5C). The differences between this map and the previous two could be explained because a higher proportion (38.4%) of the threatened taxa are already seed banked and therefore, they obtained a lower PS in the analysis. On the other hand, as only 6.3% of the potentially endemic species are represented in seed banks, they would have a higher possibility of being in the final top 100. The ability to create heatmaps for each list of species is especially advantageous for seed collection purposes, as maps may be developed based on the selected priority species.

### 4.4. Considerations for future approaches

Depending on the project goals, other resources and data sources may be incorporated into the gap analysis. At the same time, the weight of each score may be modified. For example, considering the phylogenetic diversity of the targeted species (Liu et al., 2020) or measuring the genetic variation of the collected populations (Chapman et al., 2019). Additional feedback from expert botanists and partners, as well as further revision of the literature on the biology and distribution of the species, should be considered when selecting target species for seed banking.

Fieldwork campaigns should also be planned using phenology data as periods of fruiting and dispersal are not uniform in wild species, and they can vary throughout the geographical distribution, influencing seed maturity in the potential accessions (Hay and Probert, 2013). Also, since summer and winter ephemeral plants are predicted to represent half of the species in the SD (Robichaux, 1999), collection dates should be adjusted according to local knowledge and real-time weather patterns (e.g. rainfall events). In the case of endangered species or taxa with restricted distribution, finding an adequate number of individuals with seeds could be difficult (Godefroid et al., 2011) and low-intensity, annual seed sampling from individual maternal lines will be advised to support recovery strategies.

One of the biggest challenges for the future of the plant diversity of the SD is the disparity between conservation efforts between countries. The influence of political divisions not only affects how species conservation is approached but physical walls or divisions can interfere with the connectivity and gene flow of populations (J. Liu et al., 2020; Titley et al., 2021). Planning and executing conservation projects with a transboundary approach is the only way in which the floristic diversity of the SD can be adequately preserved, especially in the context of climate change and the potential modification of the geographic distribution of species (Dallimer and Strange, 2015; López-Hoffman et al., 2010; Mason et al., 2020; Weiss and Overpeck, 2005).

Despite the limitations, seed banking continues to be one of the best approaches to protecting species (Liu et al., 2018). Best-practice integrated conservation actions (*in situ* and *ex situ*) carried out within global partnerships are essential to carrying out successful conservation projects (Breman et al., 2021). This study presents the first *ex situ* conservation gap analysis from the SD flora and provides a replicable methodology for identifying priority species and potential areas for seed bank collections.

## Acknowledgments

To the data provider institutions: California Botanic Garden, FESI-UNAM, Millennium Seed Bank, Santa Barbara Botanic Garden, San Diego Botanic Garden, San Diego Zoo Wildlife Alliance, UC Botanical Garden at Berkeley, University of California Santa Cruz, and The Arboretum at Flagstaff. To the Symbiota Support Hub for their help accessing all the coordinates from SEINet. To Juan Viruel, Itxaso Quintana, and Carolina Tovar for their valuable orientation for this project. To Armando Ponce-Vargas, Wolfgang Stuppy, and Juan Diego Arias-Montiel for the photographs in Figure 1.

## References

Alducin-Martínez, C., Ruiz Mondragón, K.Y., Jiménez-Barrón, O., Aguirre-Planter, E., Gasca-Pineda, J., Eguiarte, L.E., Medellín, R.A., 2023. Uses, Knowledge and Extinction Risk Faced by Agave Species in Mexico. Plants 12(1), 124. 10.3390/plants12010124.

Antonelli, A., Fry, C., Smith, R.J., Simmonds, M.S.J., Kersey, P.J., Pritchard, H.W., et al., 2020. State of the World”s Plants and Fungi 2020. Royal Botanic Gardens, Kew. 10.34885/172.

Bachman, S.P., Nic Lughadha, E.M., Rivers, M.C., 2018. Quantifying progress toward a conservation assessment for all plants. Conservation Biology 32, 516–524. 10.1111/cobi.13071.

Borgelt, J., Dorber, M., Høiberg, M.A., Verones, F., 2022. More than half of data deficient species predicted to be threatened by extinction. Commun Biol 5, 1–9. 10.1038/s42003-022-03638-9.

Breman, E., Ballesteros, D., Castillo-Lorenzo, E., Cockel, C., Dickie, J., Faruk, A., O”Donnell, K., Offord, C.A., Pironon, S., Sharrock, S., Ulian, T., 2021. Plant Diversity Conservation Challenges and Prospects. The Perspective of Botanic Gardens and the Millennium Seed Bank. Plants 10, 2371. 10.3390/plants10112371.

Breman, E., Balding, S., Cable, S., Carvey, N., Castillo-Lorenzo, E., Chapman, T., Cockel, C., Cossu, T.A., Dickie, J., Faruk, A., 2023. The Millennium Seed Bank Partnership: A Global Network of Seed Banks Conserving Wild Plant Species and Supporting Agriculture, Forestry, Livelihoods, and Restoration, in: Botanical Gardens and Their Role in Plant Conservation. CRC Press.

Breslin, P.B., Wojciechowski, M.F., Albuquerque, F., 2020. Projected climate change threatens significant range contraction of Cochemiea halei (Cactaceae), an island endemic, serpentine-adapted plant species at risk of extinction. Ecol Evol 10, 13211–13224. 10.1002/ece3.6914.

Brown, J.L., Cameron, A., Yoder, A.D., Vences, M., 2014. A necessarily complex model to explain the biogeography of the amphibians and reptiles of Madagascar. Nat Commun 5, 5046. 10.1038/ncomms6046.

Burke, R.A., Frey, J.K., Ganguli, A., Stoner, K.E., 2019. Species distribution modelling supports “nectar corridor” hypothesis for migratory nectarivorous bats and conservation of tropical dry forest. Diversity and Distributions 25, 1399–1415. 10.1111/ddi.12950.

Burlakova, L.E., Karatayev, A.Y., Karatayev, V.A., May, M.E., Bennett, D.L., Cook, M.J., 2011. Endemic species: Contribution to community uniqueness, effect of habitat alteration, and conservation priorities. Biological Conservation 144, 155–165. 10.1016/j.biocon.2010.08.010.

Burquez, A., Martinez-Yrizar, A., Felger, R., Yetman, D., 1999. Vegetation and habitat diversity at the southern desert edge of the Sonoran Desert. 10.2307/j.ctv34h09mn.6.

Carver, D., Sosa, C.C., Khoury, C.K., Achicanoy, H.A., Diaz, M.V., Sotelo, S., Castañeda-Álvarez, N.P., Ramirez-Villegas, J., 2021. GapAnalysis: an R package to calculate conservation indicators using spatial information. Ecography 44, 1000–1009. 10.1111/ecog.05430.

CBD, 2012. Global Strategy for Plant Conservation: 2011-2020. Botanic Gardens Conservation International, Richmond, UK.

Chapman, T., Miles, S., Trivedi, C., 2019. Capturing, protecting and restoring plant diversity in the UK: RBG Kew and the Millennium Seed Bank. Plant Diversity, Restoration of threatened plant species and their habitats 41, 124–131. 10.1016/j.pld.2018.06.001.

Corlett, R.T., 2016. Plant diversity in a changing world: Status, trends, and conservation needs. Plant Diversity 38, 10–16. 10.1016/j.pld.2016.01.001.

Cuello, W.S., Schreiber, S.J., Gremer, J.R., Venable, D.L., Trimmer, P.C., Sih, A., 2022. Extinction Risk of Sonoran Desert Annuals Following Potential Changes in Precipitation Regimes. 10.1101/2022.02.02.478887.

Dallimer, M., Strange, N., 2015. Why socio-political borders and boundaries matter in conservation. Trends in Ecology & Evolution 30, 132–139. 10.1016/j.tree.2014.12.004.

de Souza, R.A., De Marco, P., 2014. The use of species distribution models to predict the spatial distribution of deforestation in the western Brazilian Amazon. Ecological Modelling 291, 250–259. 10.1016/j.ecolmodel.2014.07.007.

Dhanjal-Adams, K.L., Quintana, I., Ballesteros, D., Carvey, N., Liu, U., Way, M., Viruel, J., Breman, E., Unpublished results. Mind the gap: global targets and strategies for ex situ conservation of Critically Endangered plants.

Diazgranados, M., Allkin, R., Black, N., Cámara-Leret, R., Canteiro, C., Carretero, J., Eastwood, R., Hargreaves, S., Hudson, A., Milliken, W., Nesbitt, M., Ondo, I., Patmore, K., Pironon, S., Turner, R., Ulian, T., Díaz-Rueda, D., 2020. World Checklist of Useful Plant Species. 10.5063/F1CV4G34.

Drezner, T.D., 2014. The keystone saguaro (Carnegiea gigantea, Cactaceae): a review of its ecology, associations, reproduction, limits, and demographics. Plant Ecol 215, 581–595. 10.1007/s11258-014-0326-y.

EPA, 2023. United States Environmental Protection Agency. Level III Ecoregions of North America. https://www.epa.gov/eco-research/ecoregions-north-america (accessed 28 March 2023).

Fick, S.E., Hijmans, R.J., 2017. WorldClim 2: new 1-km spatial resolution climate surfaces for global land areas. International Journal of Climatology 37, 4302–4315. 10.1002/joc.5086.

Girardello, M., Griggio, M., Whittingham, M.J., Rushton, S.P., 2009. Identifying important areas for butterfly conservation in Italy. Animal Conservation 12, 20–28. 10.1111/j.1469-1795.2008.00216.x.

Godefroid, S., Rivière, S., Waldren, S., Boretos, N., Eastwood, R., Vanderborght, T., 2011. To what extent are threatened European plant species conserved in seed banks? Biological Conservation, Ecoregional-scale monitoring within conservation areas, in a rapidly changing climate 144, 1494–1498. 10.1016/j.biocon.2011.01.018.

Gonzalez-Abraham, C., Garcillán, P., Ezcurra, E., Ecorregiones, G.D.T.D., 2010. Ecoregions of the Baja California peninsula: A synthesis. Boletin de la Sociedad Botanica de Mexico 87, 69–82. 10.17129/botsci.305.

Govaerts, R., Nic Lughadha, E., Black, N., Turner, R., Paton, A., 2021. The World Checklist of Vascular Plants, a continuously updated resource for exploring global plant diversity. Sci Data 8, 215. 10.1038/s41597-021-00997-6.

Hay, F.R., Probert, R.J., 2013. Advances in seed conservation of wild plant species: a review of recent research. Conservation Physiology 1, cot030. 10.1093/conphys/cot030.

Hijmans, R.J., Phillips, S., Leathwick, J., Elith, J., Hijmans, M.R.J., 2017. Package “dismo”. Circles, 9(1), 1–68.

Hijmans, R.J., Bivand, R., Forner, K., Ooms, J., Pebesma, E., Sumner, M.D., 2022. Package “terra”. Maintainer: Vienna, Austria.

Hughes, A.C., Orr, M.C., Ma, K., Costello, M.J., Waller, J., Provoost, P., Yang, Q., Zhu, C., Qiao, H., 2021. Sampling biases shape our view of the natural world. Ecography 44, 1259–1269. 10.1111/ecog.05926.

Hultine, K.R., Hernández-Hernández, T., Williams, D.G., Albeke, S.E., Tran, N., Puente, R., Larios, E., 2023. Global change impacts on cacti (Cactaceae): current threats, challenges and conservation solutions. Annals of Botany mcad040. 10.1093/aob/mcad040.

Hultine, K.R., Majure, L.C., Nixon, V.S., Arias, S., Búrquez, A., Goettsch, B., Puente-Martinez, R., Zavala-Hurtado, J.A., 2016. The Role of Botanical Gardens in the Conservation of Cactaceae. BioScience 66, 1057–1065. 10.1093/biosci/biw128.

IUCN, 2023. The IUCN Red List of Threatened Species. IUCN Red List of Threatened Species. https://www.iucnredlist.org/en (accessed 14 July 2023).

Kass, J.M., Muscarella, R., Galante, P.J., Bohl, C.L., Pinilla-Buitrago, G.E., Boria, R.A., Soley-Guardia, M., Anderson, R.P., 2021. ENMeval 2.0: Redesigned for customizable and reproducible modeling of species” niches and distributions. Methods in Ecology and Evolution 12, 1602–1608. 10.1111/2041-210X.13628

Kraus, D., Enns, A., Hebb, A., Murphy, S., Drake, D.A.R., Bennett, B., 2023. Prioritizing nationally endemic species for conservation. Conservation Science and Practice 5, e12845. 10.1111/csp2.12845.

Leroy, B., Meynard, C.N., Bellard, C., Courchamp, F., 2016. virtualspecies, an R package to generate virtual species distributions. Ecography 39(6), 599–607. 10.1111/ecog.01388.

Li, D.-Z., Pritchard, H.W., 2009. The science and economics of ex situ plant conservation. Trends Plant Sci 14, 614–621. 10.1016/j.tplants.2009.09.005.

Liu, J., Yong, D.L., Choi, C.-Y., Gibson, L., 2020. Transboundary Frontiers: An Emerging Priority for Biodiversity Conservation. Trends in Ecology & Evolution 35, 679–690. 10.1016/j.tree.2020.03.004.

Liu, U., Breman, E., Cossu, T.A., Kenney, S., 2018. The conservation value of germplasm stored at the Millennium Seed Bank, Royal Botanic Gardens, Kew, UK. Biodivers Conserv 27, 1347–1386. 10.1007/s10531-018-1497-y.

Liu, U., Cossu, T.A., Davies, R.M., Forest, F., Dickie, J.B., Breman, E., 2020. Conserving orthodox seeds of globally threatened plants ex situ in the Millennium Seed Bank, Royal Botanic Gardens, Kew, UK: the status of seed collections. Biodivers Conserv 29, 2901–2949. 10.1007/s10531-020-02005-6.

Liu, U., Gianella, M., Aranda, P., Diazgranados, M., Flores-Ortiz, C., Lira, R., Bacci, S., Mattana, E., Milliken, W., Mitrovits, O., Pritchard, H., Rodríguez-Arévalo, I., Way, M., Williams, C., Ulian, T., 2023. Conserving useful plants for a sustainable future: species coverage, spatial distribution, and conservation status within the Millennium Seed Bank collection. Biodiversity and Conservation 32. 10.1007/s10531-023-02631-w.

López-Hoffman, L., Varady, R.G., Flessa, K.W., Balvanera, P., 2010. Ecosystem services across borders: a framework for transboundary conservation policy. Frontiers in Ecology and the Environment 8, 84–91. 10.1890/070216.

Margulies, J.D., Moorman, F.R., Goettsch, B., Axmacher, J.C., Hinsley, A., 2022. Prevalence and perspectives of illegal trade in cacti and succulent plants in the collector community. Conservation Biology 37, e14030. 10.1111/cobi.14030.

Martyn-Yenson, A., Offord, C.A., Meagher, P.F., Auld, T.D., Bush, D., Coates, D.J., Commander, L.E., Guja, L.K., Norton, S., Makinson, R.O., Stanley, R., Walsh, N., Wrigley, D., Broadhurst, L., 2021. Plant Germplasm Conservation in Australia: strategies and guidelines for developing, managing and utilising ex situ collections. Third edition. Australian Network for Plant Conservation, Canberra.

Mason, N., Ward, M., Watson, J.E.M., Venter, O., Runting, R.K., 2020. Global opportunities and challenges for transboundary conservation. Nat Ecol Evol 4, 694–701. 10.1038/s41559-020-1160-3.

McLaughlin, S.P., Bowers, J.E., 1999. Diversity and Affinities of the Flora of the Sonoran Floristic Province, in: Ecology of Sonoran Desert Plants and Plant Communities. University of Arizona Press, pp. 12–25.

Merritt, D.J., Dixon, K.W., 2011. Restoration Seed Banks. A Matter of Scale. Science 332, 424–425. 10.1126/science.1203083.

NatureServe, 2023. About the Data. NatureServe Explorer. https://explorer.natureserve.org/AboutTheData (accessed 14 July 2023).

Ortega-Baes, P., Godínez-Alvarez, H., 2006. Global Diversity and Conservation Priorities in the Cactaceae. Biodivers Conserv 15, 817–827. 10.1007/s10531-004-1461-x.

Pammenter, N.W., Berjak, P., 2013. Development of the understanding of seed recalcitrant and implications for ex situ conservation. Biotecnología Vegetal 13, 131–144.

Parsons, E.C.M., 2016. Why IUCN Should Replace “Data Deficient” Conservation Status with a Precautionary “Assume Threatened” Status—A Cetacean Case Study. Frontiers in Marine Science 3. 10.3389/fmars.2016.00193.

Pebesma, E.J., 2018. Simple features for R: standardized support for spatial vector data. R J., 10(1), 439–446.

Pence, V.C., 2013. In Vitro Methods and the Challenge of Exceptional Species for Target 8 of the Global Strategy for Plant Conservation1. mobt 99, 214–220. 10.3417/2011112.

Pence, V.C., Ballesteros, D., Walters, C., Reed, B.M., Philpott, M., Dixon, K.W., Pritchard, H.W., Culley, T.M., Vanhove, A.-C., 2020. Cryobiotechnologies: Tools for expanding long-term ex situ conservation to all plant species. Biological Conservation 250, 108736. 10.1016/j.biocon.2020.108736.

Phillips, S.J., Anderson, R.P., Schapire, R.E., 2006. Maximum entropy modeling of species geographic distributions. Ecological Modelling 190, 231–259. 10.1016/j.ecolmodel.2005.03.026

Pironon, S., Ondo, I., Diazgranados, M., Allkin, R., Baquero, A.C., Cámara-Leret, R., Canteiro, C., Dennehy-Carr, Z., Govaerts, R., Hargreaves, S., Hudson, A.J., Lemmens, R., Milliken, W., Nesbitt, M., Patmore, K., Schmelzer, G., Turner, R.M., van Andel, T.R., Ulian, T., Antonelli, A., Willis, K.J., 2024. The global distribution of plants used by humans. Science 383, 293–297. 10.1126/science.adg8028

R Core Team, 2023. R: A language and environment for statistical computing. R Foundation for Statistical Computing, Vienna, Austria. URL https://www.R-project.org/.

Radosavljevic, A., Anderson, R.P., 2014. Making better Maxent models of species distributions: complexity, overfitting and evaluation. Journal of biogeography, 41(4), 629–643. 10.1111/jbi.12227.

Ramírez-Villegas, J., Khoury, C., Jarvis, A., Debouck, D.G., Guarino, L., 2010. A Gap Analysis Methodology for Collecting Crop Genepools: A Case Study with Phaseolus Beans. PLOS ONE 5, e13497. 10.1371/journal.pone.0013497.

RBG Kew, 2023. Seed Collection, Millennium Seed Bank. https://www.kew.org/science/collections-and-resources/collections/seed-collection (accessed 14 July 2023).

Rebman, J., Roberts, C., 2012. Baja California Plant Field Guide. San Diego, CA.

Riemann, H., Ezcurra, E., 2005. Plant endemism and natural protected areas in the peninsula of Baja California, Mexico. Biological Conservation 122, 141–150. 10.1016/j.biocon.2004.07.008.

Rivière, S., Müller, J.V., 2018. Contribution of seed banks across Europe towards the 2020 Global Strategy for Plant Conservation targets, assessed through the ENSCONET database. Oryx 52, 464–470. 10.1017/S0030605316001496.

Robichaux, R.H., 1999. Ecology of Sonoran Desert Plants and Plant Communities. University of Arizona Press.

Rowe, H., Montijo, A.B., 2020. Sonoran Desert Plant Specialist Group. https://www.mcdowellsonoran.org/wp-content/uploads/2021/09/121-2020-Sonoran-Desert-Plant-SG-Report_RFP.pdf (accessed 14 April 2023).

Scott, J.M., Davis, F., Csuti, B., Noss, R., Butterfield, B., Groves, C., Anderson, H., Caicco, S., D”Erchia, F., Edwards, T.C., Ulliman, J., Wright, R.G., 1993. Gap Analysis: A Geographic Approach to Protection of Biological Diversity. Wildlife Monographs 3–41.

SEINet, 2023. SEINet Portal Network. http://swbiodiversity.org/seinet/index.php. (accessed 24 May 2023).

SEMARNAT, 2010. NORMA OFICIAL MEXICANA NOM-059-SEMARNAT-2010. Procuraduría Federal de Protección al Ambiente. https://www.gob.mx/profepa/documentos/norma-oficial-mexicana-nom-059-semarnat-2010 (accessed 14 July 2023).

Shreve, F., Wiggins, I.L., 1951. Vegetation and Flora of the Sonoran Desert: Vegetation of the Sonoran Desert, by F. Shreve. Carnegie Institution of Washington.

Sowa, S.P., Annis, G., Morey, M.E., Diamond, D.D., 2007. A Gap Analysis and Comprehensive Conservation Strategy for Riverine Ecosystems of Missouri. Ecological Monographs 77, 301–334. 10.1890/06-1253.1.

Teixido, A.L., Toorop, P.E., Liu, U., Ribeiro, G.V.T., Fuzessy, L.F., Guerra, T.J., Silveira, F.A.O., 2017. Gaps in seed banking are compromising the GSPC”s Target 8 in a megadiverse country. Biodivers Conserv 26, 703–716. 10.1007/s10531-016-1267-7.

Tinoco-Ojanguren, C., Reyes-Ortega, I., Sánchez-Coronado, M.E., Molina-Freaner, F., Orozco-Segovia, A., 2016. Germination of an invasive Cenchrus ciliaris L. (buffel grass) population of the Sonoran Desert under various environmental conditions. South African Journal of Botany 104, 112–117. 10.1016/j.sajb.2015.10.009.

Titley, M.A., Butchart, S.H.M., Jones, V.R., Whittingham, M.J., Willis, S.G., 2021. Global inequities and political borders challenge nature conservation under climate change. Proceedings of the National Academy of Sciences 118, e2011204118. 10.1073/pnas.2011204118.

Tovar, C., Hudson, L., Cuesta, F., Meneses, R.I., Muriel, P., Hidalgo, O., Palazzesi, L., Suarez Ballesteros, C., Hammond Hunt, E., Diazgranados, M., Hind, D.J.N., Forest, F., Halloy, S., Aguirre, N., Baker, W.J., Beck, S., Carilla, J., Eguiguren, P., Françoso, E., Gámez, L.E., Jaramillo, R., Llambí, L.D., Maurin, O., Melcher, I., Muller, G., Roy, S., Viñas, P., Yager, K., Viruel, J., 2023. Strategies of diaspore dispersal investment in Compositae: the case of the Andean highlands. Annals of Botany mcad099. 10.1093/aob/mcad099.

Trejo-Salazar, R.-E., Eguiarte, L.E., Suro-Piñera, D., Medellin, R.A., 2016. Save Our Bats, Save Our Tequila: Industry and Science Join Forces to Help Bats and Agaves. naar 36, 523–530. 10.3375/043.036.0417.

Turner, R.M., Brown, D.E., 1982. Sonoran Desertscrub. CALS Publications Archive. The University of Arizona.

Tweddle, J.C., Dickie, J.B., Baskin, C.C., Baskin, J.M., 2003. Ecological aspects of seed desiccation sensitivity. Journal of Ecology 91, 294–304. 10.1046/j.1365-2745.2003.00760.x

UNEP-WCMC & IUCN, 2023. Protected Planet: The World Database on Protected Areas (WDPA) and World Database on Other Effective Area-based Conservation Measures (WD-OECM). http://www.protectedplanet.net (accessed 14 July 2023).

van Slageren, W., 2003. The Millennium Seed Bank: building partnerships in arid regions for the conservation of wild species. Journal of Arid Environments 54, 195–201. 10.1006/jare.2001.0879.

Velazco, S.J.E., Rose, M.B., de Andrade, A.F.A., Minoli, I., Franklin, J., 2022. flexsdm: An r package for supporting a comprehensive and flexible species distribution modelling workflow. Methods in Ecology and Evolution, 13(8), 1661–1669. 10.1111/2041-210X.13874.

Vignali, S., Barras, A.G., Arlettaz, R., Braunisch, V., 2020. SDMtune: An R package to tune and evaluate species distribution models. Ecology and Evolution, 10(20), 11488–11506. 10.1002/ece3.6786.

Way, M., 2003. Chapter 9 Collecting Seed from Non-domesticated Plants for Long-Term Conservation. Seed Conservation: Turning Science into Practice.

Wei, T., Simko, V., Levy, M., Xie, Y., Jin, Y., Zemla, J., 2017. Package “corrplot”. Statistician, 56(316), e24.

Weiss, J., Overpeck, J., 2005. Is the Sonoran Desert losing its cool? Global Change Biology 11, 2065–2077. 10.1111/j.1365-2486.2005.01020.x.

Wickham, H., Averick, M., Bryan, J., Chang, W., McGowan, L. D. A., François, R., et al., 2019. Welcome to the Tidyverse. Journal of open source software 4(43), 1686. 10.21105/joss.01686.

Wiggins, I.L., 1980. Flora of Baja California. Flora of Baja California.

Wilson, M.F., Leigh, L., Felger, R.S., 2002. Invasive exotic plants in the Sonoran Desert. Invasive exotic species in the Sonoran region.

Wyse, S.V., Dickie, J.B., 2018. Taxonomic affinity, habitat and seed mass strongly predict seed desiccation response: a boosted regression trees analysis based on 17 539 species. Annals of Botany 121, 71–83. 10.1093/aob/mcx128.

Wyse, S.V., Dickie, J.B., 2017. Predicting the global incidence of seed desiccation sensitivity. Journal of Ecology 105, 1082–1093. 10.1111/1365-2745.12725

Ye, J., Shan, Z., Peng, D., Sun, M., Niu, Y., Liu, Y., Zhang, Q., Yang, Y., Lin, Q., Chen, J., Zhu, R., Wang, Y., Chen, Z., 2023. Identifying gaps in the ex situ conservation of native plant diversity in China. Biological Conservation 282, 110044. 10.1016/j.biocon.2023.110044.

Yin, Y., He, Q., Pan, X., Liu, Q., Wu, Y., Li, X., 2022. Predicting Current Potential Distribution and the Range Dynamics of Pomacea canaliculata in China under Global Climate Change. Biology 11, 110. 10.3390/biology11010110.

