## Supplementary material for "Sonoran Desert *ex situ* conservation gap analysis: charting the path towards conservation"

List of the Top 100 taxa selected by the species prioritisation.

| Ranking | Family | Species | Author | Priority Score |
|---|---|---|---|---|
| 1 | Asparagaceae | *Agave pelona* | Gentry | 0.775 |
| 2 | Asparagaceae | *Agave turneri* | R.H.Webb & Salazar-Ceseña | 0.719 |
| 3 | Apocynaceae | *Vallesia laciniata* | Brandegee | 0.717 |
| 4 | Asparagaceae | *Agave zebra* | Gentry | 0.700 |
| 5 | Cactaceae | *Cochemiea boolii* | (G.E.Linds.) P.B.Breslin & Majure | 0.645 |
| 6 | Nyctaginaceae | *Mirabilis greenei* | S.Watson | 0.637 |
| 7 | Asteraceae | *Encelia ravenii* | Wiggins | 0.625 |
| 8 | Fabaceae | *Lupinus arizonicus subsp. Arizonicus* | (S. Watson) S. Watson | 0.625 |
| 9 | Rutaceae | *Thamnosma trifoliata* | I.M.Johnst. | 0.625 |
| 10 | Rubiaceae | *Galium mechudoense* | Dempster | 0.625 |
| 11 | Cactaceae | *Grusonia marenae* | (S.H.Parsons) E.F.Anderson | 0.624 |
| 12 | Lamiaceae | *Salvia palmetorum* | J.G.González & Carnahan | 0.624 |
| 13 | Verbenaceae | *Citharexylum shrevei* | Moldenke | 0.624 |
| 14 | Rubiaceae | *Stenaria sanchezii* | Lorence | 0.623 |
| 15 | Malpighiaceae | *Malpighia watsonii* | Rose | 0.583 |
| 16 | Asteraceae | *Verbesina felgeri* | B.L.Turner | 0.562 |
| 17 | Asparagaceae | *Agave azurea* | R.H.Webb & G.D.Starr | 0.562 |
| 18 | Cactaceae | × *Cylindronia robertsii* | (Rebman) M.A.Baker, Majure, Cloud-H. & Rebman | 0.562 |
| 19 | Cucurbitaceae | *Cucurbita cylindrata* | L.H.Bailey | 0.537 |
| 20 | Asparagaceae | *Agave gigantensis* | Gentry | 0.525 |
| 21 | Rubiaceae | *Galium moranii* | Dempster | 0.520 |
| 22 | Asparagaceae | *Agave vizcainoensis* | Gentry | 0.520 |
| 23 | Cactaceae | *Cochemiea angelensis* | (R.T.Craig) P.B.Breslin & Majure | 0.520 |

| | | | | |
|---|---|---|---|---|
| 24 | Cactaceae | *Cochemiea cerralboa* | (Britton & Rose) P.B.Breslin & Majure | 0.520 |
| 25 | Meliaceae | *Swietenia humilis* | Zucc. | 0.506 |
| 26 | Cucurbitaceae | *Echinopepon minimus var. minimus* | (Kellogg) S.Watson | 0.500 |
| 27 | Malvaceae | *Allobriquetia sonorae* | (Fryxell) Bovini | 0.500 |
| 28 | Cactaceae | *Ferocactus emoryi subsp. covillei* | (Britton & Rose) D.R.Hunt & Dimmitt | 0.500 |
| 29 | Anacardiaceae | *Rhus kearneyi subsp. kearneyi* | F.A. Barkley | 0.500 |
| 30 | Solanaceae | *Lycium californicum var. arizonicum* | A.Gray | 0.500 |
| 31 | Crassulaceae | *Dudleya cymosa subsp. cymosa* | (Lem.) Britton & Rose | 0.500 |
| 32 | Fabaceae | *Astragalus newberryi var. aquarii* | Isely | 0.500 |
| 33 | Rubiaceae | *Stenotis asperuloides var. brandegeeana* | (Rose) Terrell | 0.500 |
| 34 | Asteraceae | *Coreocarpus sonoranus var. libranus* | B.L.Turner | 0.500 |
| 35 | Cactaceae | *Pelecyphora alversonii* | (J.M.Coult.) D.Aquino & Dan.Sánchez | 0.500 |
| 36 | Polygonaceae | *Chorizanthe rosulenta* | Reveal | 0.500 |
| 37 | Boraginaceae | *Phacelia pauciflora* | S.Watson | 0.500 |
| 38 | Acanthaceae | *Ruellia comonduensis* | T.F.Daniel | 0.500 |
| 39 | Lamiaceae | *Hedeoma tenuiflora* | Brandegee | 0.500 |
| 40 | Lamiaceae | *Monardella thymifolia* | Greene | 0.500 |
| 41 | Polygonaceae | *Eriogonum pilosum* | S.Stokes | 0.500 |
| 42 | Asteraceae | *Acourtia palmeri* | (S.Watson) Reveal & R.M.King | 0.500 |
| 43 | Verbenaceae | *Verbena calinfera* | G.L.Nesom | 0.500 |
| 44 | Nyctaginaceae | *Mirabilis oligantha* | (Standl.) J.F.Macbr. | 0.500 |
| 45 | Rubiaceae | *Galium volcanense* | Dempster | 0.500 |
| 46 | Asteraceae | *Verbesina palmeri* | S.Watson | 0.500 |
| 47 | Asteraceae | *Senecio pinacatensis* | Felger | 0.500 |
| 48 | Plantaginaceae | *Penstemon vizcainensis* | Moran | 0.500 |
| 49 | Crassulaceae | *Dudleya linearis* | (Greene) Britton & Rose | 0.500 |
| 50 | Crassulaceae | *Dudleya pachyphytum* | Moran & M.Benedict | 0.500 |
| 51 | Polemoniaceae | *Linanthus viscainensis* | Moran | 0.500 |
| 52 | Polygonaceae | *Chorizanthe mutabilis* | Brandegee | 0.500 |

| | | | | |
|---|---|---|---|---|
| 53 | Asteraceae | *Heterotheca thiniicola* | (Rzed. & E.Ezcurra) B.L.Turner | 0.500 |
| 54 | Boraginaceae | *Cryptantha pondii* | Greene | 0.500 |
| 55 | Polygonaceae | *Chorizanthe flava* | Brandegee | 0.500 |
| 56 | Crassulaceae | *Dudleya rubens* | (Brandegee) Britton & Rose | 0.500 |
| 57 | Fabaceae | *Astragalus orcuttianus* | S.Watson | 0.500 |
| 58 | Amaranthaceae | *Amaranthus tucsonensis* | Henrickson | 0.500 |
| 59 | Rubiaceae | *Spermacoce lagunensis* | (M.E.Jones) Govaerts | 0.500 |
| 60 | Asteraceae | *Verbesina oligocephala* | I.M.Johnst. | 0.500 |
| 61 | Asteraceae | *Laphamia ajoensis* | (Todsen) Lichter-Marck | 0.500 |
| 62 | Euphorbiaceae | *Euphorbia platysperma* | Engelm. | 0.500 |
| 63 | Solanaceae | *Datura arenicola* | Gentry ex Bye & Luna | 0.500 |
| 64 | Polygonaceae | *Eriogonum moranii* | Reveal | 0.500 |
| 65 | Solanaceae | *Lycium densifolium* | Wiggins | 0.500 |
| 66 | Polygonaceae | *Eriogonum preclarum* | Reveal | 0.500 |
| 67 | Polygonaceae | *Eriogonum intricatum* | Benth. | 0.500 |
| 68 | Polygonaceae | *Eriogonum angelense* | Moran | 0.500 |
| 69 | Fabaceae | *Hoffmannseggia peninsularis* | (Britton) Wiggins | 0.499 |
| 70 | Asteraceae | *Perityle carterae* | (A.M.Powell) Lichter-Marck | 0.499 |
| 71 | Asteraceae | *Bajacalia moranii* | B.L.Turner | 0.499 |
| 72 | Polemoniaceae | *Bryantiella palmeri* | (S.Watson) J.M.Porter | 0.499 |
| 73 | Cistaceae | *Crocanthemum nutans* | (Brandegee) Janch. | 0.499 |
| 74 | Asteraceae | *Hofmeisteria filifolia* | I.M.Johnst. | 0.499 |
| 75 | Campanulaceae | *Nemacladus australis* | (Munz) Morin | 0.499 |
| 76 | Cactaceae | *Cochemiea thornberi* | (Orcutt) P.B.Breslin & Majure | 0.499 |
| 77 | Brassicaceae | *Dimorphocarpa pinnatifida* | Rollins | 0.499 |
| 78 | Cactaceae | *Echinocereus grandis* | Britton & Rose | 0.499 |
| 79 | Polemoniaceae | *Dayia scabra* | (Brandegee) J.M.Porter | 0.499 |
| 80 | Brassicaceae | *Lyrocarpa linearifolia* | Rollins | 0.499 |
| 81 | Plantaginaceae | *Mecardonia exilis* | (Brandegee) Pennell | 0.499 |

| | | | | |
|---|---|---|---|---|
| 82 | Polemoniaceae | *Dayia grantii* | J.M.Porter | 0.499 |
| 83 | Poaceae | *Distichlis bajaensis* | H.L.Bell | 0.499 |
| 84 | Fabaceae | *Acmispon nudatus* | (Greene) Brouillet | 0.499 |
| 85 | Fabaceae | *Acmispon flexuosus* | (Greene) Brouillet | 0.499 |
| 86 | Plantaginaceae | *Sairocarpus virga* | (A.Gray) D.A.Sutton | 0.499 |
| 87 | Verbenaceae | *Citharexylum roxanae* | Moldenke | 0.499 |
| 88 | Asteraceae | *Hofmeisteria anomalochaeta* | (R.M.King) B.L.Turner | 0.499 |
| 89 | Cactaceae | *Cylindropuntia ciribe* | (Engelm. ex J.M.Coult.) F.M.Knuth | 0.498 |
| 90 | Cactaceae | *Cochemiea estebanensis* | (G.E.Linds.) P.B.Breslin & Majure | 0.498 |
| 91 | Martyniaceae | *Martynia palmeri* | S.Watson | 0.498 |
| 92 | Cactaceae | *Cylindropuntia libertadensis* | Rebman | 0.498 |
| 93 | Cactaceae | *Lophophora williamsii* | (Lem. ex J.F.Cels) J.M.Coult. | 0.496 |
| 94 | Zygophyllaceae | *Viscainoa pinnata* | (I.M.Johnst.) Gentry | 0.494 |
| 95 | Olacaceae | *Ximenia glauca* | (DeFilipps) Bentouil | 0.493 |
| 96 | Cucurbitaceae | *Cucurbita cordata* | S.Watson | 0.488 |
| 97 | Euphorbiaceae | *Acalypha saxicola* | Wiggins | 0.485 |
| 98 | Cactaceae | *Opuntia ficus-indica* | (L.) Mill. | 0.475 |
| 99 | Malvaceae | *Hibiscus tiliaceus* | (Guill. & Perr.) Steud. | 0.475 |
| 100 | Euphorbiaceae | *Bernardia gentryana* | Croizat | 0.466 |
